# Visualizing whole-head neuronal activity and information-sharing dynamics in *Caenorhabditis elegans* during the transition into the anesthetized state

**DOI:** 10.64898/2026.03.31.715586

**Authors:** Hamilton White, Cameron R. Bosinski, Hortense Bogaert, Christopher V. Gabel, Christopher W. Connor

**Affiliations:** Department of Anesthesiology, Mass General Brigham, Boston, MA, USA; Harvard Medical School, Harvard University, Boston, MA, USA; Department of Pharmacology, Physiology and Biophysics, Chobanian & Avedisian School of Medicine, Boston University, Boston, MA, USA; Department of Biology, Northeastern University, Boston, MA, USA

## Abstract

General anesthetics, such as isoflurane, induce immobility and decrease responsiveness to external stimuli. Our previous studies used *C. elegans* to investigate recovery from anesthesia, and established information-theory metrics characterizing how system-wide neuronal dynamics differ under these conditions. This study develops a novel technique enabling the corollary experiment: visualization of neuronal activity throughout the transition into the anesthetized state. *C. elegans* worms expressing pan-neuronal nuclear RFP and cytosolic GCaMP6s were imaged with wide-field cameras (behavior), and a light sheet microscope (neuronal activity) in most neurons in the animal’s head during isoflurane exposure. Stable concentrations were maintained by measured flow vaporization of isoflurane into a specially designed gas enclosure compatible with the imaging system. We analyzed ensemble neuronal activity, spectrograms of frequency over time, and metrics of shared information between neurons. Behavioral decline differed among tested isoflurane conditions and was most pronounced at 8% isoflurane. Spectrograms indicate loss of bulk signal power, with a somewhat greater loss in higher frequency bands. State Decoupling, Internal Predictability, and System Consistency were among the useful metrics for discriminating anesthetized state. The ratio of State Decoupling to System Consistency (“Disconnection Ratio”) showed a prominent increase over time for the 8% condition. The induction of anesthesia produces progressive changes in neuronal activity and communication that can be quantified at single-neuron resolution. Information metrics (State Decoupling entropy, System Consistency entropy, and their ratio) demonstrate that the isoflurane anesthetization process results in altered entropy-derived information sharing, particularly at high anesthetic concentrations.

## Introduction

Despite their ubiquity, mechanisms by which volatile anesthetics alter neural function are poorly understood [1]. Many mechanistic hypotheses include targets such as ryanodine receptors, agonizing inhibitory GABA signaling, and mitochondrial effects [2–8]. Abrupt changes in EEG frequency power have been observed upon anesthetic induction, but whether this reflects single cell dynamics is underexplored [9]. *Caenorhabditis elegans* is a powerful model for performing this analysis at single-cell resolution due to its compact nervous system, translucent body, tractable genetics, and rapid growth [10,11].

Previously, we tracked 120 neurons in the *C. elegans* head and quantified their activity via functional imaging of cytoplasmic calcium concentrations using fluorescent reporters [12]. Under the influence of the anesthetic, there is a loss of positively correlated neuronal activity and a breakdown of neuronal signaling as measured by information shared between neurons [10,11]. We have further shown that anesthesia-mediated neural activity changes can be identified using novel entropy-based metrics, such as state decoupling and internal predictability. While analyzing *C. elegans* during removal from anesthesia, we observed convergence of these metrics with isoflurane-free populations over a period of ∼2 hrs [10].

However, our previous experiments have not assessed neuronal activity during the continuous transition from the awake to the anesthetized state. In this work, we accomplish two discrete tasks: 1) We create a microscope-compatible gas enclosure allowing continuous recording *C. elegans* behavior and nervous system activity during continuous isoflurane exposure. 2) Given this recording system, we assess how gross animal immobility and neuronal kinetics change throughout the onset of anesthesia. Details of our past microscopy techniques, experimental design, fluorescence images and examples of extracted neuron activity are illustrated in Fig. 1.

**Figure 1:**
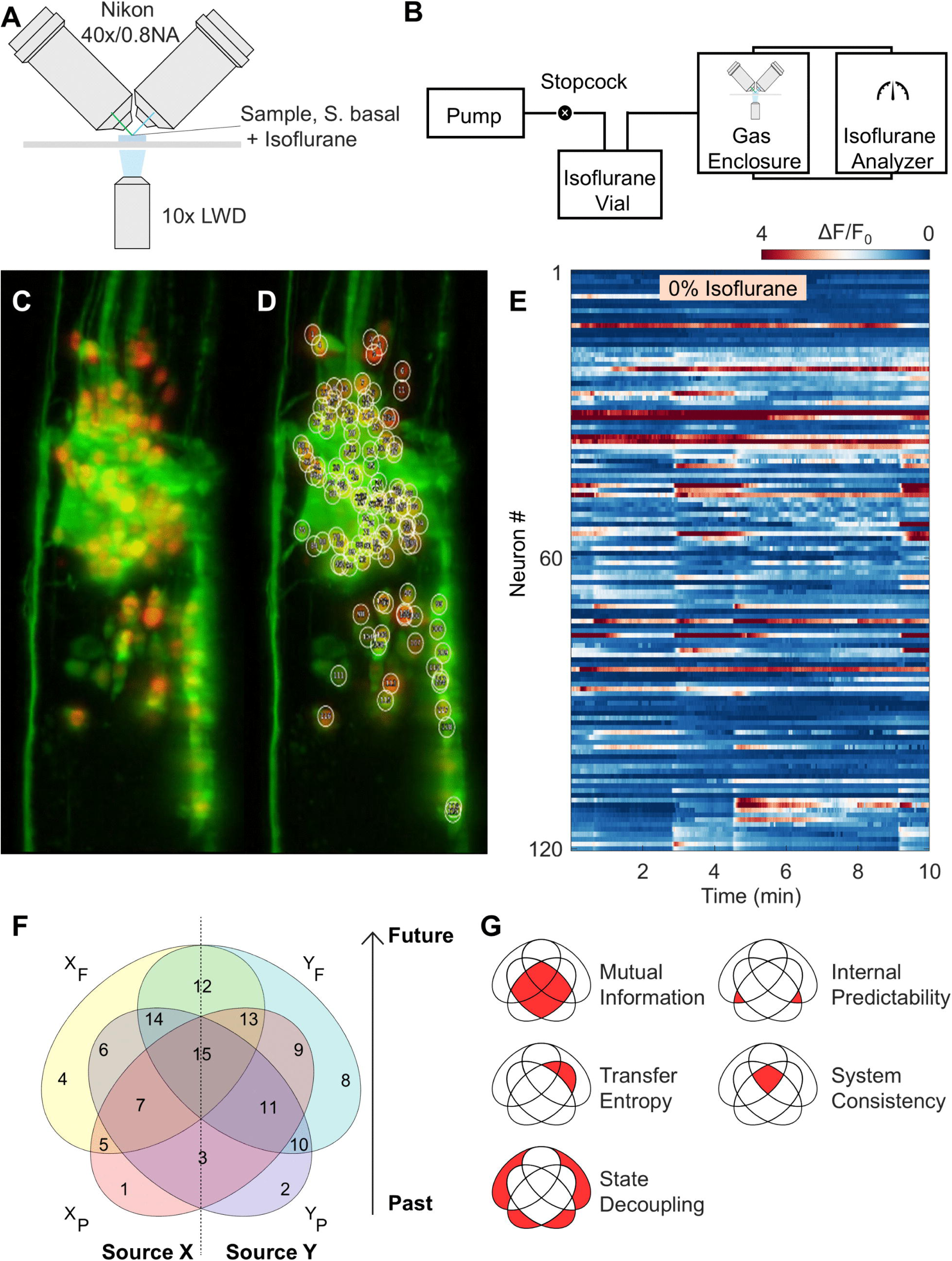
Tracking of *Caenorhabditis elegans* neuronal activity and computing functional entropy metrics. Diagrams in F and G correspond with, and are adapted from, definitions in our prior work *Chang et al (2023)*. (A) Example diagram of a Dual-view Inverted Selective Plane Illumination Microscope (diSPIM) as used for capturing pan-neuronal activity in *C. elegans*. (B) Anesthetic delivery apparatus, which delivers a constant flow of anesthetic vapor to a closed circuit composed of an anesthetic enclosure and an infrared spectroscopy gas analyzer. (C) Example image of *C. elegans* animal recorded using diSPIM microscopy. This view is the result of taking a maximum projection of the first recorded imaging volume. Red puncta indicate neuronal nuclei. Green shows the relative position of the cytoplasmic GCaMP genetically encoded calcium sensor. (D) Tracking summary for an animal recorded in (A-C). White circles indicate the automatically identified nuclei. Numbering within each circle indicates the neuron ID # assigned to each respective nucleus identified, and the corresponding row of the activity matrix. (E) Summary raster heatmap of normalized activity from animal recorded and tracked in (D) for a total of 10 minutes duration. An isoflurane condition of 0% is shown. (F) Venn diagram of overlapping regions between two sources (X & Y) and two timepoints (past [P] and future [F]). Summation of entropy percentage computed in each region produces a measure of the corresponding amount of information shared between each source and timepoint. (G) Definition of five key entropy metrics from *Chang et al (2023).* Each metric is the summation of entropy for the regions of the diagram in (F), with the same definitions as our prior publication. Metrics computed for this work include Mutual Information, Transfer Entropy, State Decoupling, Internal Predictability, and System Consistency.

## Materials and Methods

### C. elegans Strains

Experiments were performed on young adult hermaphrodites (3 days after egg-laying) of strain QW1144;QW1155 *(zfis125[rgef-1p::GCaMP6s], otIs355[rab-3p::NLS::tagRFP])*. Both GCaMP6s (a fluorescent calcium reporter) and nuclear-localized RFP are expressed pan-neuronally in this strain (gift of Mark Alkema, University of Massachusetts). *C. elegans* were cultivated at 20°C per standard protocols [13]. Animals were chosen at random and were handled in the same manner across experimental days. While *C. elegans* males do exist, their low prevalence (0.2-0.5%), and structurally different nervous system make them infeasible for this work [14,15].

### Pan-neuronal Imaging using diSPIM Light-Sheet Microscopy

Light-sheet microscopy was performed to capture spontaneous *in vivo* neural activity within 120 neurons in the animal’s head region at single-cell resolution, using a protocol previously described in detail across several publications [10,12,16,17]. Briefly, animals were first immobilized in 20 μL droplets of 5 mM L-levamisole in S-basal buffer, then encapsulated in 8 µL polyethylene glycol diacrylate (PEGDA, a highly biocompatible polymer) hydrogel disks bound to #1.5 coverslips, bathed in the paralytic buffer for imaging. One hour of time under levamisole elapsed before imaging initiation. Imaging used a dual-inverted selective plane illumination fluorescence microscope (schematic in Fig. 1A, ASI) with water immersion 40×/0.8NA objectives (Nikon USA, Melville, NY), and 10 mW excitation intensity. Each animal was imaged at two volumes/second (XYZ: 0.1625×0.1625×1 μm, 7320 timepoints recorded), capturing RFP and GCaMP6s fluorescence. Each animal was exposed to the isoflurane protocol once, then discarded.

Isoflurane concentrations of 0%, 2% and 8% were employed, approximately corresponding to 0, 0.6, and 2.6 MAC-like concentrations respectively in *C. elegans* [18]. 2% and 8% (mild and very high doses) reflect the technically achievable bounds of the dosage space for this preparation [11,19]. Sample sizes were selected to match our prior study using the same method [10]. Animals exposed to no isoflurane (0%) served as immobilized, isoflurane-free controls. We expected exposure to isoflurane at 2% (∼0.6 MAC) in *C. elegans* to produce subtle differences in neuronal activity, whereas exposure to 8% isoflurane (∼2.6 MAC) should induce a profound loss of activity.

### Isoflurane Delivery

A custom gas enclosure was placed around the dish containing encapsulated *C. elegans* and the microscope objectives. Three small-bore catheters were introduced: for vapor delivery, gas sampling by infrared spectroscopy, and gas analyzer sample return. The gas sampling line and sample return line form a closed circuit (Fig. 1B). Isoflurane concentrations in the chamber could be continuously sampled by the gas analyzer and a known concentration returned to the enclosure. Separately, compressed air was directed over a reservoir of isoflurane to deliver isoflurane vapor into the enclosure, rate-adjusted to maintain the desired concentration over time. Low flow rates were adequate to compensate for small system leaks (with initial rise to steady state in approximately one minute). Recording of neural activity started concurrently with initiation of isoflurane flow. Continuous vigilance is required to titrate isoflurane into the experimental apparatus during imaging, hence experimenters cannot be blinded to the experimental condition. Further data reduction was automated, and we analyzed all acquired data that met our quality-of-imaging standards (i.e., neurons recorded were in focus, remained in view throughout acquisition).

### Behavioral Quantification

A widefield camera (ASI120MM Mini, ZWO), telescoping C-mount optics, and AstroDMx software allowed recording swimming animals in 1 mL of 1x S. basal buffer for one hour at 0%, 2%, and 8% isoflurane [13]. We believed one hour to be sufficient to achieve equilibration of gaseous isoflurane with the liquid environment. Sixty seconds of images were recorded at a rate of 10 fps, one recording every 5 minutes (13 total videos). Images were analyzed on a high performance computer to determine swimming rates per animal (characterizing anesthetic behavioral response) using custom implementations of *wrMTrck* and *pyimagej* [20,21].

### Computational Analysis of Neuronal Imaging

Custom Python scripts were employed during post processing to track the brightest 120 in-frame RFP-labelled nuclei per animal notated anatomically from the tip of the animal’s head, in three dimensions. GCaMP6s signals were extracted from the surrounding somas as previously described (Fig. 1C-E) [11]. The first 120 timepoints of all datasets were cropped to remove a period of intense photobleaching present at the beginning of all pan-neuronal recordings, rendering a total analyzed recording length of 7200 timepoints (60 minutes). We normalized activity as ΔF/F_0_ with F_0_ defined as the bottom first percentile of each neuron’s activity level per trace. Spectrograms were computed from the first derivative of normalized neuronal activity traces using Chartrand’s algorithm [22]. FFT spectra were computed per-neuron in 60 one-minute epochs, then averaged to create each part of the spectrogram. Interpolative shading smoothed the visualization. Individual power spectra were computed after removing the DC component from the recording. The Power Spectral Density was used to quantify individual frequency components of the global responses observed [23]. Loss of signal power was computed using percentage loss from the first computed timepoint (minute one), with bounds of the mean per concentration (of minute one) as 100% and -150 dB as the 0%, which is 15 dB below the lowest computed timepoint signal power of -134.8 dB. Entropy metrics were generated using mathematical functions described in previous work, and the definitions of those metrics were reused exactly here (Fig. 1F/G) [10,24]. Entropy measures were smoothed per-animal using a moving mean method and a window size of 40 computed timepoints.

### Statistics

Swimming behavior was quantified as body bends per minute (BBPM). For each acquisition set and time point, mean BBPM was calculated from all tracked segments within the corresponding timepoint and acquisition. Acquisition set was treated as the biological replicate for statistical analysis: total biological replicates obtained were 0% n=6, 2% n=5, 8% n=5. The total number of tracked *C. elegans* per isoflurane condition and timepoint were >150, which in aggregate exceeds similar automated analyses from previous work [20,25]. For experimental consistency, at least one 0%, 2%, and 8% experiment was done each experimental day. The analysis of behavioral trajectories used a linear mixed-effects model with fixed effects of isoflurane concentration, time, and their interaction. Time was centered at 30 minutes to assess the impact of isoflurane concentration on the rate of immobilization. Estimates were obtained using restricted maximum likelihood (REML), and significance testing used Satterthwaite approximated degrees of freedom [26].

For entropy metrics, pairwise comparisons used a Kruskal-Wallis test with Bonferroni correction. Adjusted p<0.05 was used for statistical significance. Statistical values were kept exact to three decimal places and compiled or exported automatically to Microsoft Excel spreadsheet files (see S1-S6 Files). All metrics were computed in MATLAB (MathWorks, Natick, MA).

## Results

### Behavioral recording reveals a concentration-dependent reduction in mobility for animals exposed to isoflurane

*C. elegans* exposed to atmospheres of constant isoflurane concentrations show a progressive reduction in mobility over time as assessed by their swimming behavior. *C. elegans* swim with rhythmic sinusoidal motion when placed into a liquid environment, and Body Bends Per Minute (BBPM) is a well-established and automatable measurement of this form of mobility, thus enabling rapid screening [13,20,27–29]. It takes time for isoflurane to diffuse through the liquid column and equilibrate into the bodies of the *C. elegans*. Thus, animals swam extensively during early exposure to the isoflurane atmosphere (Fig. 2A) but, after one hour of equilibration time, we observed a marked decrease in mobility (Fig. 2B).

**Figure 2:**
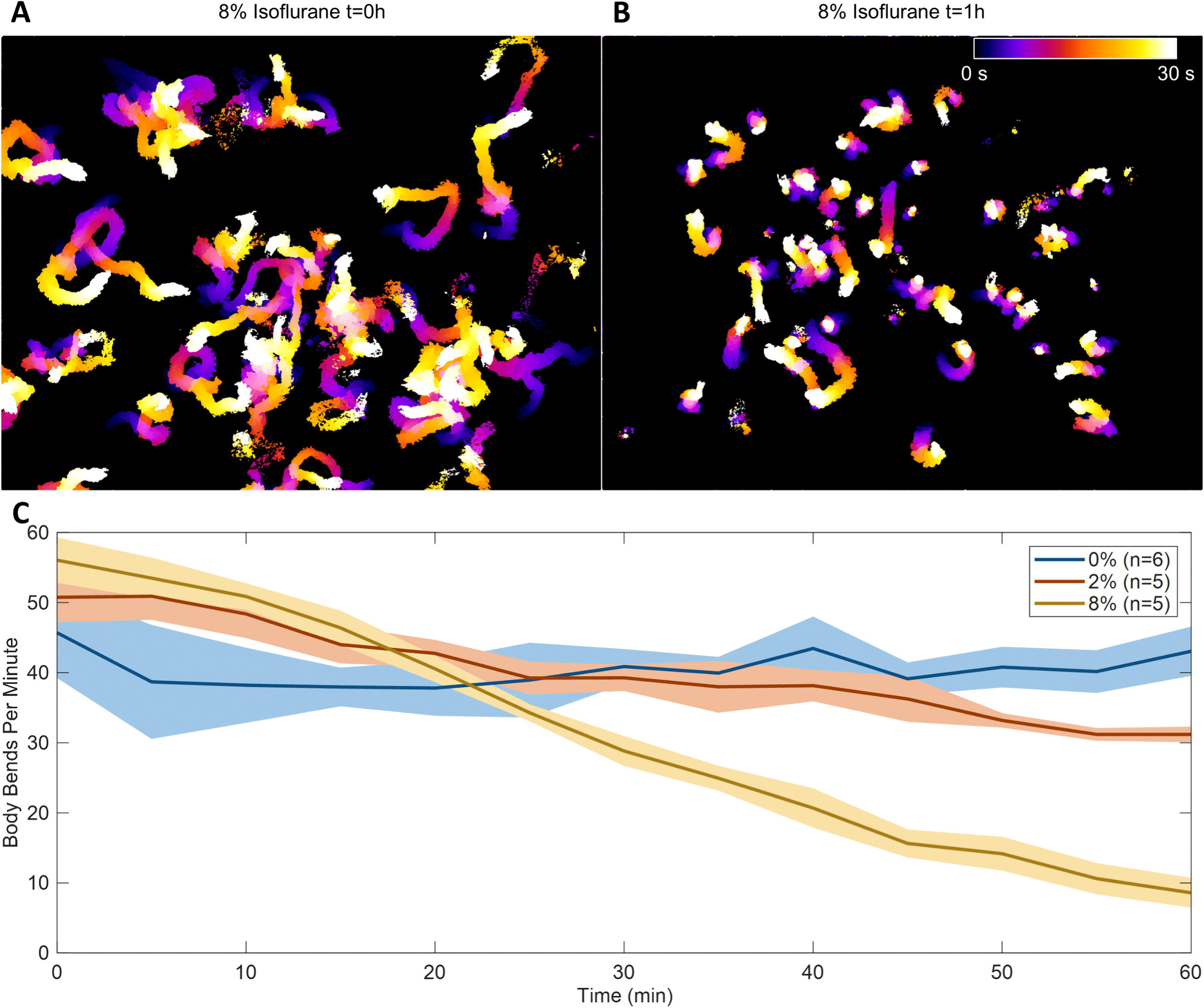
Behavioral analysis of animal movement during continuous isoflurane exposure. Behavioral measurements were obtained from a separate, non-immobilized cohort. (A) Colored timelapse overlay of the extent of movement for example animals exposed to an 8% isoflurane atmosphere at the beginning of the recording sequence. (B) Colored timelapse overlay of the same overall population of animals in (A) after 1 hour exposure. (C) Time-course plot of mean swimming movement for 0%, 2%, and 8% isoflurane conditions. Shaded regions indicate 95% confidence intervals around the mean. Biological replicates of 0%: 6, 2%: 5, and 8%: 5 were recorded. In aggregate >150 tracks were included in each isoflurane condition and timepoint.

Mean population BBPM was analyzed at the acquisition-set level (mean traces per condition are shown in Fig. 2C). A random-slope model fit the data better than a random-intercept-only model (likelihood-ratio test, Χ^2^=60.23, p<0.001). The concentration-by-time interaction was significant (F(2,12.7)=21.15, p<0.001): higher isoflurane concentrations do produce greater amounts of immobilization over time. Estimated swimming activity remained stable in the 0% condition (+0.025 BBPM/min), declined under 2% isoflurane (-0.34 BBPM/min), and declined rapidly to 8% isoflurane conditions (-0.89 BBPM/min). The slope difference between 8% and 2% remained significant after Bonferroni correction (Δ=-0.52 BBPM/min, simultaneous 95% CI [-0.87, -0.17], adjusted p=0.009), whereas the difference between 2% and 0% did not reach significance after correction (Δ=-0.37 BBPM/min, simultaneous 95% CI [-0.87, -0.17], adjusted p=0.059).

### Pan-neuronal recording during isoflurane exposure allows visualization of the neuronal kinetics of anesthetic induction

Figure 3A shows examples of normalized heatmaps for real-time recordings of neuronal activity in 120 individual neurons in the head of *C. elegans* during exposure to 0%, 2%, and 8% over 60 minutes. *C. elegans* are observed to become behaviorally immobilized over this time-period (Fig. 2), with effects at 8% being stronger than 2% [1,18]. Qualitatively, the heatmaps in Fig. 3A show more loss of activity with higher concentrations of isoflurane.

**Figure 3:**
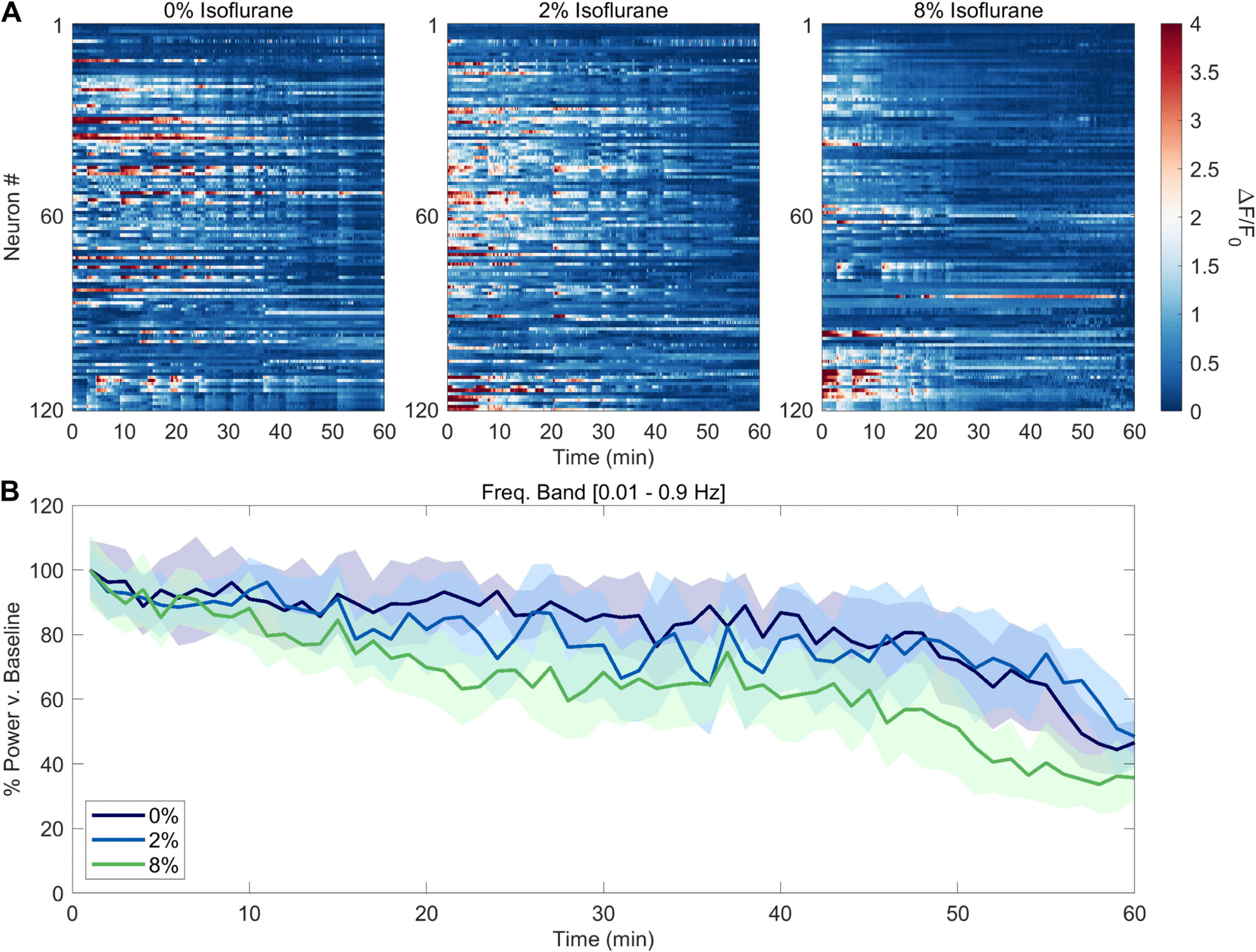
Neural activity during continuous exposure to isoflurane. (A) Example heatmaps of normalized neural activity for 120 neurons at 0%, 2%, and 8% isoflurane conditions. Each example is similarly color scaled. (B) Time-course plot of mean power relative to the first retained analysis window (the first minute of recorded data) for neural activity of the recorded population of animals at 0%, 2%, and 8% isoflurane concentration (sample sizes n=8).

In human studies, EEG spectrograms display the evolution in the power contribution of component signal frequencies over time. This characterizes the changes in the architecture of the frontal EEG during the initiation of general anesthesia, but the signal arises from the combined activity of millions of neurons per EEG lead [30,31]. Our recordings presented the opportunity to compute similar power spectra over the individual neurons acquired (up to a Nyquist limit of 1 Hz due to the sampling rate of 2 fps), building a spectrogram over each minute of neuronal imaging per animal. The bulk power dropped significantly more for the 8% condition than the 0% or 2% condition (Figure 3B). The exact kinetics vary substantially per animal (Fig. 4), but secondary analyses of discrete frequency bands (0.01–0.1 Hz, 0.1-0.5 Hz, and 0.5-1 Hz, see S1 Fig.) showed some indication of band-specific differences in loss of power between concentrations after t=50 minutes.

**Figure 4:**
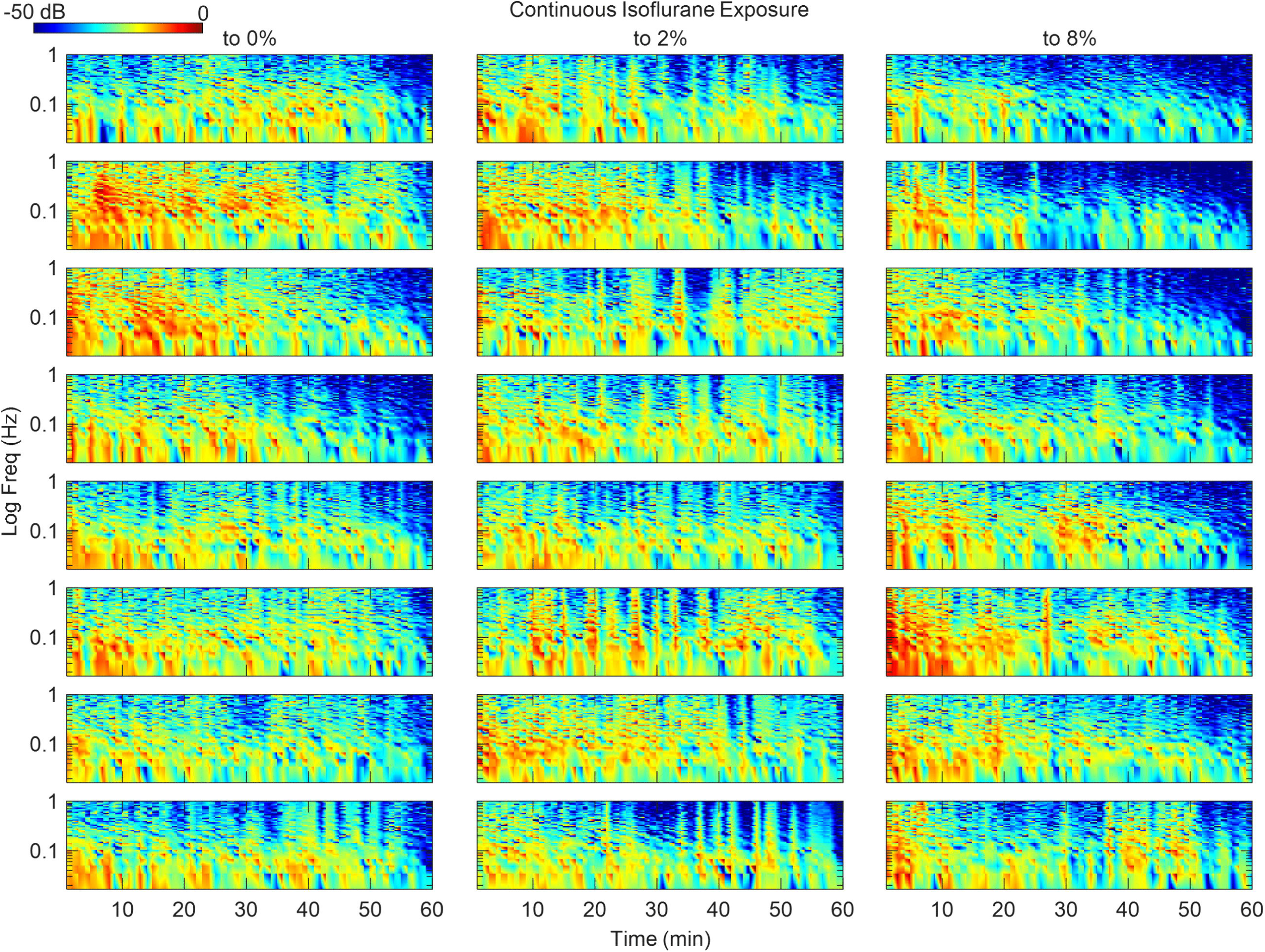
Real time interpolated spectrograms for individual animals exposed to continuous isoflurane exposure at stable 0, 2, and 8% conditions. Spectrograms were computed on a per-minute basis for the entire recording duration of 60 minutes. Y-axis scales are log frequencies in hertz. Scale bar on the upper left indicates signal power (dB). Sample sizes of n=8 per condition were analyzed (down each column). Spectral plots were color-scaled equivalently.

### Evolution of information theory metrics during the transition from the awake to the anesthetized state

To measure the degree of connectivity and organization in the system, we employed entropy metrics that quantify the amount of information shared between neurons and over time, as previously described [10]. At each timepoint, for each pairing of two neurons (X & Y), we compute the entropy of their adjacent past (i.e., X_P_ and Y_P_) and future states (e.g., X_F_ and Y_F_), and hence the entropy metrics shown in Fig. 1G. The evolution of these metrics is shown in Fig. 5.

**Figure 5:**
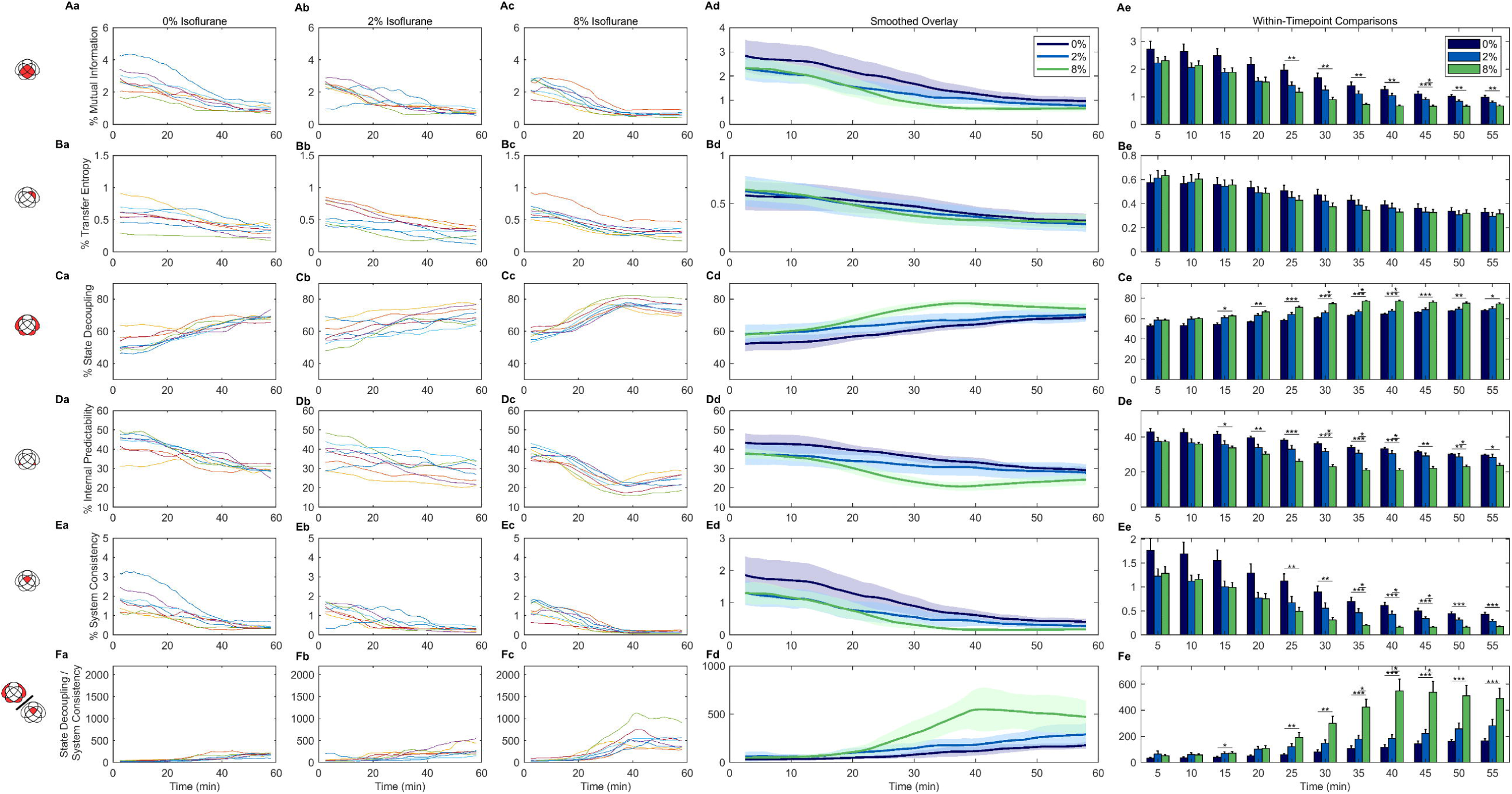
Matrix of time-course evolution for entropy parameters of individual animals and a summary metric of system disconnection. (A-F) Each row plots a different entropy metric in the following order: % Mutual Information, % Transfer Entropy, % State Decoupling, % Internal Predictability, % System Consistency, and Disconnection Ratio (% State Decoupling / % System Consistency). Information in each column displays the same category of information plotted for each overall entropy metric in (A-F). (a) Per-animal entropy traces at 0% isoflurane concentration. (b) Per-animal entropy traces at 2% isoflurane concentration. (c) Per-animal entropy traces at 8% isoflurane concentration. (d) Smoothed overlay of interval of traces by concentration in (a-c). Shaded regions indicate 95% confidence interval. (e) Statistical comparison of discrete timepoint data every 5 minutes for the 5-55 minute range of data in column (d). Error bars indicate SEM. * p<0.05, ** p<0.01, *** p<0.001. Kruskal-Wallis, Bonferroni correction for comparisons of 0%, 2%, and 8% within a given timepoint. Exact multiple comparisons in column (e) are shown in S1-S6 Files. Sample sizes of n=8 per isoflurane condition. See S1-S6 Files for exact pairwise comparisons, 95% confidence intervals, and p-values.

Individual entropy metrics moved in a temporally mirrored manner to our previous results regarding anesthetic removal (Fig. 5A,B,C,D,E, S1-S6 Files) [10]. The most differences between discrete timepoints observed were within a time-range of 30-50 minutes after initiation. Mutual Information between neurons and timepoints (i.e., quantifying the information shared between two neurons over time) decreased progressively with increasing isoflurane concentration (Fig. 5A). While transfer entropy (i.e., the information specifically transferred from one neuron to the future of its paired neuron) did decrease at higher anesthetic concentrations, this reflected only a small proportion of the total entropy and was not statistically significantly different (Fig. 5B,Be). The State Decoupling (i.e., the amount of system information/entropy not shared between neurons or timepoints) of animals induced with 8% isoflurane increased significantly from the 0% and 2% conditions (Fig. 5C), before plateauing after t=40 minutes. Conversely, Internal Predictability (i.e., how much an individual neuron’s activity is consistent over time) demonstrated a reciprocal relationship with State Decoupling (Fig. 5D). System Consistency (i.e., how much information is shared across the system over time) showed progressive decline to small values over time (Fig. 5E). Lastly, we computed the ratio of State Decoupling and System Consistency, to use as a metric of how disconnected the system was becoming (“Disconnection Ratio,” Fig. 5(f), rightmost column). Discrete timepoint comparisons at 5-minute intervals of Disconnection Ratio (Kruskal-Wallis with Bonferroni correction) found that there were more significant differences across isoflurane concentrations between t=30 minutes and t=50 minutes (see S1-S6 Files for exact within-timepoint pairwise comparisons).

To verify the robustness of these analyses, we asked whether creating an alternate within-neuron disconnection ratio (state decoupling divided by internal predictability) would net the same trends in concentration and time. It does (S2A Fig.). We further tested the robustness of the results through reducing the computational step-size and reducing the number of quantization levels to just two. The shape of the disconnection ratio curves did not change from our other computed methods (S2B Fig.). Lastly, we removed the DC drift signal caused by photobleaching using per-neuron normalization of the neuronal activity trace, finding that the trends in Fig. 5 remained observable (S2C Fig.).

## Discussion

General anesthesia is canonically understood as a clinically-inducible brain state characterized by loss of normal behavior and drastic changes in neuronal activity [32]. Previous publications of our pan-neuronal imaging modality showed a reduction of shared information upon step-wise presentation of increasing isoflurane concentrations, and that continuous imaging after anesthetic removal showed a distinct reversal of this reduction concurrent with a resumption of neuronal activity dynamics [10,11,18]. We asked: would the kinetics of transitioning into the anesthetized state resemble our previous observations during anesthetic removal? We created a gas enclosure to deliver atmospheres of consistent isoflurane concentrations during longitudinal mobility recording. Using swimming as a coarse behavioral metric, we found that there was a significant reduction in mobility during anesthetic exposure (Fig. 2). Zero percent isoflurane conditions were flat, 2% showed a modest reduction in mobility, while 8% showed the highest amount of mobility reduction. We found loss of mobility to be smooth and progressive.

Pan-neuronal imaging using light sheet microscopy allowed the real-time analysis of neuronal activity, with a greater loss of neural activity at higher isoflurane concentrations (Fig. 3). This trend holds across frequency bands, though lower frequency activity was less distinguishable than higher frequency across concentrations (S1 Fig). The exact timing of spectrographic effects was individualized per animal (Fig. 4). These outcomes follow logically both from our previous work and studies in larger mammalian models showing a dose-dependent increase in attenuation at higher frequencies [10,11,33,34].

In prior work, we developed five entropy metrics derived fundamentally from Shannon Entropy, and evaluated how those metrics (Mutual Information, Transfer Entropy, State Decoupling, Internal Predictability, and System Consistency) evolve with time as an animal recovers from anesthesia [10,35,36]. Here, we applied the same approach, assessing anesthetic exposure. We found that State Decoupling entropy (the amount of information temporally segregated within a given neuron) increases with increasing isoflurane concentration (Fig. 5). Internal Predictability entropy (the amount of shared information within a neuron over time) moves in reciprocity to state decoupling (Fig. 5). System Consistency decreased in the system over time across conditions (the amount of information shared across time between neurons). Across all metrics, statistical differences were greatest between 8% and 0%/2% for timepoints recorded between t=30 and t=50 minutes, and the overall trend mirrors our previous study on removal kinetics.

Since State Decoupling and System Consistency reflect key aspects of global temporal dynamics, we combined them into a descriptive composite metric (Disconnection Ratio) to summarize differences between local and global organization. If this disconnection ratio increases, there is less shared information between neurons and timepoints. Control populations demonstrated a flat trend in disconnection ratio over time (Fig. 5). Animals exposed to 8% isoflurane show a strong increase in the ratio (Fig. 5). To address concerns of robustness, replacing System Consistency with Internal Predictability yielded a qualitatively similar result, suggesting choice of denominator is not determinative (S2A Fig.). Other computational techniques evaluated yielded comparable results (S2B,C Fig.). These results conceptually match findings from our prior work, larger mammals, and human data [18,37–39]. The Disconnection Ratio reaches a maximum before flattening out over time; this may reflect a saturation of the metric under the substantial suppression of neuronal activity observed. The neuronal dynamics after this timepoint represent a highly suppressed system.

The work presented in this paper has limitations. Behavioral measurements originated from freely swimming animals, whereas animals immobilized with levamisole and PEGDA encapsulation were necessary for pan-neuronal imaging. Although all neuronal imaging groups had identical handling, interactions between levamisole and isoflurane exposure cannot be excluded. Thus, we cannot determine the within-animal temporal relationship between behavioral responsiveness and neuronal dynamics. Also, the study examined exposures to atmospheres of 0%, 2%, and 8% isoflurane; intermediate concentrations were not evaluated. Isoflurane delivery began concurrently with imaging acquisition. The first minute of recording was removed because of photobleaching. Consequently, the first retained timepoint represents an early-exposure measurement rather than a pre-exposure baseline. Although the entropy metrics remained robust across multiple analytical approaches, generalized suppression of neuronal dynamics may still confound results.

Overall, our work establishes a new method for exposing *C. elegans* to volatile anesthetics while continuously recording gross behavior or neuronal activity at single cell resolution. In future work, we will ask what features of the communication circuit make an animal differentially sensitive to anesthetics, whether these results hold across genetic backgrounds, and to what extent other anesthetic agents produce different effects on the segregation of information.

## Acknowledgements

We acknowledge Andrew Chang, Daniel Taub, and Belen Karakullukcu for their advice in the preparation of this work. We also acknowledge the Boston University Neurophotonics Training Program.

## Supporting Information

**S1 Fig: Spectrogram data in finite frequency bands.** Time-course plot of mean power relative to the first retained analysis timepoint for neural activity in related figure 3. Data was separated into discrete frequency bands and plotted separately. A) Mean power versus initial timepoint for frequency band [0.01, 0.1] Hz. B) Plot as in (A) for frequency band [0.1, 0.5] Hz. C) Plots as in (A/B) for frequency band [0.5, 1] Hz. Shaded regions indicate 95% confidence intervals.

**S2 Fig: Alternative disconnection ratio and robustness analyses.** A) Alternative normalization for disconnection ratio as in Figure 5Fd (0%, 2%, and 8% isoflurane concentrations). Here the ratio quantities are % State Decoupling and % Internal Predictability. B) Robustness analysis of entropy calculations using smaller windows for entropy calculations (50% of that displayed in Figure 5), and only two discrete quantization levels for neural activity versus the four discrete states used in Figure 5. C) Robustness analysis of entropy calculations using the ‘normalize’ function in MATLAB to remove the DC drift signal by neuron prior to four-level quantization, as in Figure 5 and Figure S3A. Shaded area around each line is the 95% confidence interval.

**S1 File: Within-timepoint statistics related to Fig. 5Ae**.

**S2 File: Within-timepoint statistics related to Fig. 5Be**.

**S3 File: Within-timepoint statistics related to Fig. 5Ce**.

**S4 File: Within-timepoint statistics related to Fig. 5De**.

**S5 File: Within-timepoint statistics related to Fig. 5Ee**.

**S6 File: Within-timepoint statistics related to Fig. 5Fe**.

## References

1. Kelz MB, Mashour GA. The Biology of General Anesthesia from Paramecium to Primate. Curr Biol. 2019;29: R1199–R1210. doi:10.1016/j.cub.2019.09.071

2. Kanaya HJ, Kuwajima K, Ito Y, Shinohara Y, Okubo Y, Shiono S, et al. Isoflurane activates the type 1 ryanodine receptor to induce anesthesia in mice. PLoS Biol. 2025;23: e3003172. doi:10.1371/journal.pbio.3003172

3. Topf N, Jenkins A, Baron N, Harrison NL. Effects of isoflurane on gamma-aminobutyric acid type A receptors activated by full and partial agonists. Anesthesiology. 2003;98: 306–311. doi:10.1097/00000542-200302000-00007

4. Jia F, Yue M, Chandra D, Homanics GE, Goldstein PA, Harrison NL. Isoflurane is a potent modulator of extrasynaptic GABA(A) receptors in the thalamus. J Pharmacol Exp Ther. 2008;324: 1127–1135. doi:10.1124/jpet.107.134569

5. Hentschke H, Raz A, Krause BM, Murphy CA, Banks MI. Disruption of cortical network activity by the general anaesthetic isoflurane. Br J Anaesth. 2017;119: 685–696. doi:10.1093/bja/aex199

6. Bukhari Q, Schroeter A, Rudin M. Increasing isoflurane dose reduces homotopic correlation and functional segregation of brain networks in mice as revealed by resting-state fMRI. Sci Rep. 2018;8: 10591. doi:10.1038/s41598-018-28766-3

7. Kayser E-B, Morgan PG, Sedensky MM. GAS-1 : A Mitochondrial Protein Controls Sensitivity to Volatile Anesthetics in the Nematode Caenorhabditis elegans. Anesthesiology. 1999;90: 545. doi:10.1097/00000542-199902000-00031

8. Morgan PG, Kayser E-B, Sedensky MM. C. elegans and volatile anesthetics. WormBook: The Online Review of C elegans Biology [Internet]. WormBook; 2007. Available: https://www.ncbi.nlm.nih.gov/books/NBK19697/

9. Jiang J, Zhao Y, Liu J, Yang Y, Liang P, Huang H, et al. Signatures of Thalamocortical Alpha Oscillations and Synchronization With Increased Anesthetic Depths Under Isoflurane. Front Pharmacol. 2022;13: 887981. doi:10.3389/fphar.2022.887981

10. Chang AS, Wirak GS, Li D, Gabel CV, Connor CW. Measures of information content during anesthesia and emergence in the C. elegans nervous system. Anesthesiology. 2023;139: 49–62. doi:10.1097/ALN.0000000000004579

11. Awal MR, Wirak GS, Gabel CV, Connor CW. The collapse of global neuronal states in C. elegans under isoflurane anesthesia. Anesthesiology. 2020;133: 133–144. doi:10.1097/ALN.0000000000003304

12. Wirak GS, Florman J, Alkema MJ, Connor CW, Gabel CV. Age-associated changes to neuronal dynamics involve a disruption of excitatory/inhibitory balance in C. elegans. Zimmer M, Sengupta P, editors. eLife. 2022;11: e72135. doi:10.7554/eLife.72135

13. Stiernagle T. Maintenance of C. elegans. WormBook. 2006 [cited 28 Apr 2020]. doi:10.1895/wormbook.1.101.1

14. Brenner S. The Genetics of CAENORHABDITIS ELEGANS. Genetics. 1974;77: 71–94.

15. Walsh JD, Boivin O, Barr MM. What About the Males? The C. elegans Sexually Dimorphic Nervous System and a CRISPR-Based Tool to Study Males in a Hermaphroditic Species. J Neurogenet. 2020;34: 323–334. doi:10.1080/01677063.2020.1789978

16. Burnett K, Edsinger E, Albrecht DR. Rapid and gentle hydrogel encapsulation of living organisms enables long-term microscopy over multiple hours. Commun Biol. 2018;1. Available: https://www.ncbi.nlm.nih.gov/pmc/articles/PMC6123791/

17. Han X, Su Y, White H, O’Neill KM, Morgan NY, Christensen R, et al. A polymer index-matched to water enables diverse applications in fluorescence microscopy. Lab Chip. 2021;21: 1549–1562.

18. Awal M, Austin D, Florman J, Alkema M, Gabel CV, Connor CW. The breakdown of neural function under isoflurane anesthesia: in vivo, multineuronal imaging in C. elegans. Anesthesiology. 2018;129: 733–743. doi:10.1097/ALN.0000000000002342

19. Chang AS, Mazuera L, Gabel CV, Connor CW. Anesthesia isn’t sleep: The neuronal dynamics of immobility in isoflurane-anesthetized C. elegans differ from the activity patterns of previously established sleep-like quiescent states. PLoS One. 2025;20: e0324323. doi:10.1371/journal.pone.0324323

20. Nussbaum-Krammer CI, Neto MF, Brielmann RM, Pedersen JS, Morimoto RI. Investigating the spreading and toxicity of prion-like proteins using the metazoan model organism C. elegans. J Vis Exp. 2015; 52321. doi:10.3791/52321

21. Rueden CT, Hiner MC, Evans EL, Pinkert MA, Lucas AM, Carpenter AE, et al. PyImageJ: A library for integrating ImageJ and Python. Nat Methods. 2022;19: 1326–1327. doi:10.1038/s41592-022-01655-4

22. Chartrand R. Numerical Differentiation of Noisy, Nonsmooth Data. International Scholarly Research Notices. 2011;2011: 164564. doi:10.5402/2011/164564

23. Gao R. Interpreting the electrophysiological power spectrum. J Neurophysiol. 2016;115: 628–630. doi:10.1152/jn.00722.2015

24. Ting HK. On the Amount of Information. Theory of Probability & Its Applications. 1962;7: 439–447. doi:10.1137/1107041

25. Albrecht DR, Bargmann CI. High-content behavioral analysis of Caenorhabditis elegans in precise spatiotemporal chemical environments. Nat Methods. 2011;8: 599–605. doi:10.1038/nmeth.1630

26. Heise MJ, Mon SK, Bowman LC. Utility of linear mixed effects models for event-related potential research with infants and children. Developmental Cognitive Neuroscience. 2022;54: 101070. doi:10.1016/j.dcn.2022.101070

27. Buckingham SD, Sattelle DB. Fast, automated measurement of nematode swimming (thrashing) without morphometry. BMC Neurosci. 2009;10: 84. doi:10.1186/1471-2202-10-84

28. Hahm J-H, Kim S, DiLoreto R, Shi C, Lee S-JV, Murphy CT, et al. C. elegans maximum velocity correlates with healthspan and is maintained in worms with an insulin receptor mutation. Nat Commun. 2015;6: 8919. doi:10.1038/ncomms9919

29. Currey HN, Liachko NF. Evaluation of Motor Impairment in <EM>C. elegans</EM< Models of Amyotrophic Lateral Sclerosis. JoVE (Journal of Visualized Experiments). 2021; e62699. doi:10.3791/62699

30. Connor CW. Open Reimplementation of the BIS Algorithms for Depth of Anesthesia. Anesth Analg. 2022; 10.1213/ANE.0000000000006119. doi:10.1213/ANE.0000000000006119

31. Purdon PL, Sampson A, Pavone KJ, Brown EN. Clinical Electroencephalography for Anesthesiologists: Part I: Background and Basic Signatures. Anesthesiology. 2015;123: 937. doi:10.1097/ALN.0000000000000841

32. Brown EN, Lydic R, Schiff ND. General Anesthesia, Sleep, and Coma. N Engl J Med. 2010;363: 2638–2650. doi:10.1056/NEJMra0808281

33. Hudetz AG, Vizuete JA, Pillay S. Differential Effects of Isoflurane on High-Frequency and Low-Frequency γ Oscillations in the Cerebral Cortex and Hippocampus in Freely Moving Rats. Anesthesiology. 2011;114: 588–595. doi:10.1097/ALN.0b013e31820ad3f9

34. Kortelainen J, Jia X, Seppänen T, Thakor N. Increased electroencephalographic gamma activity reveals awakening from isoflurane anaesthesia in rats. Br J Anaesth. 2012;109: 782–789. doi:10.1093/bja/aes265

35. Shannon CE. A mathematical theory of communication. The Bell System Technical Journal. 1948;27: 379–423. doi:10.1002/j.1538-7305.1948.tb01338.x

36. Schreiber T. Measuring Information Transfer. Phys Rev Lett. 2000;85: 461–464. doi:10.1103/PhysRevLett.85.461

37. Luppi AI, Golkowski D, Ranft A, Ilg R, Jordan D, Bzdok D, et al. General anaesthesia decreases the uniqueness of brain functional connectivity across individuals and species. Nat Hum Behav. 2025; 1–18. doi:10.1038/s41562-025-02121-9

38. Cohen D, Swinderen B van, Tsuchiya N. Isoflurane Impairs Low-Frequency Feedback but Leaves High-Frequency Feedforward Connectivity Intact in the Fly Brain. eNeuro. 2018;5. doi:10.1523/ENEURO.0329-17.2018

39. Gelbard-Sagiv H, Mudrik L, Hill MR, Koch C, Fried I. Human single neuron activity precedes emergence of conscious perception. Nat Commun. 2018;9: 2057. doi:10.1038/s41467-018-03749-0

