## Supplementary figures and images for "Visualizing whole-head neuronal activity and information-sharing dynamics in *Caenorhabditis elegans* during the transition into the anesthetized state"

### Supplemental Fig 1

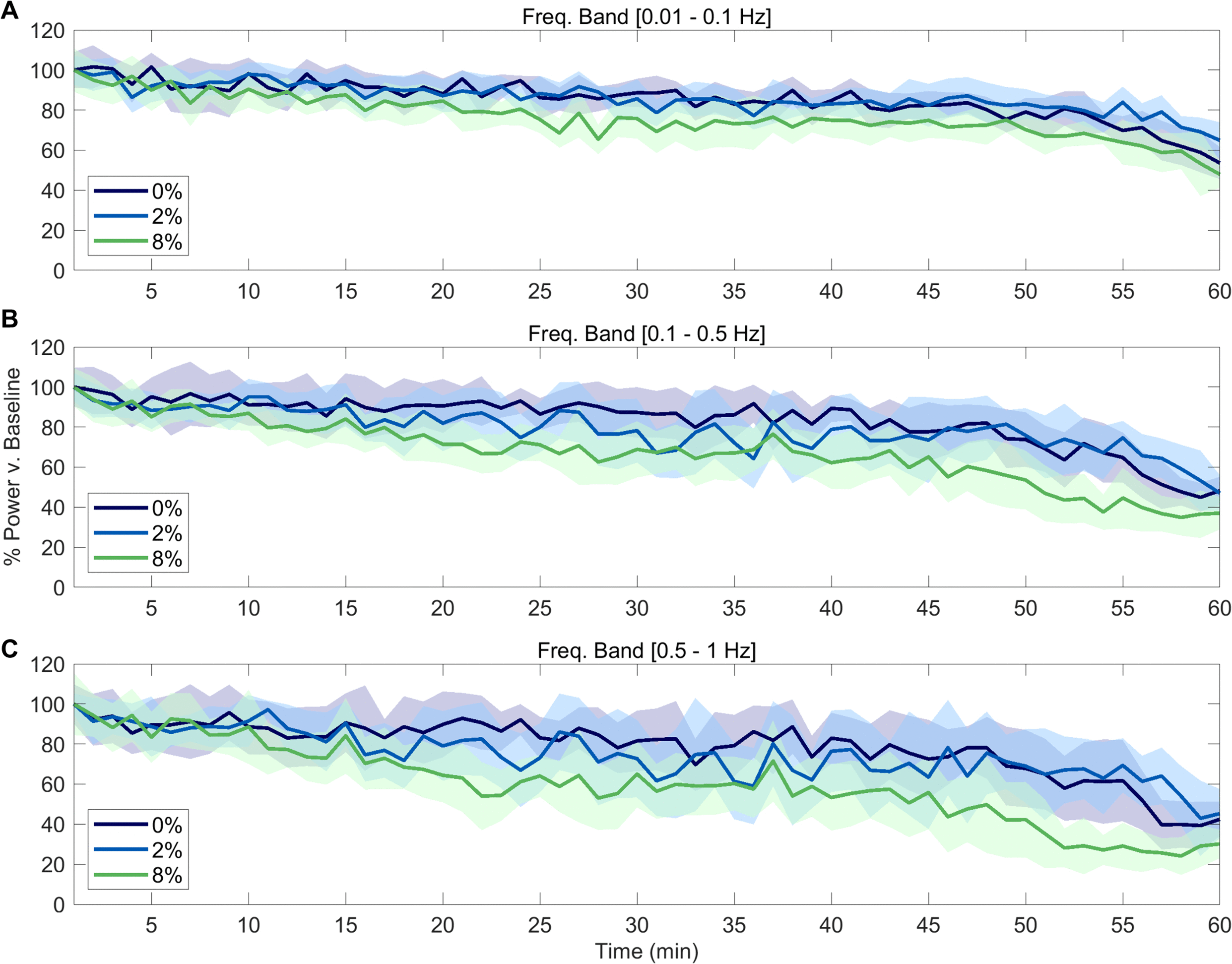

### Supplemental Fig 2

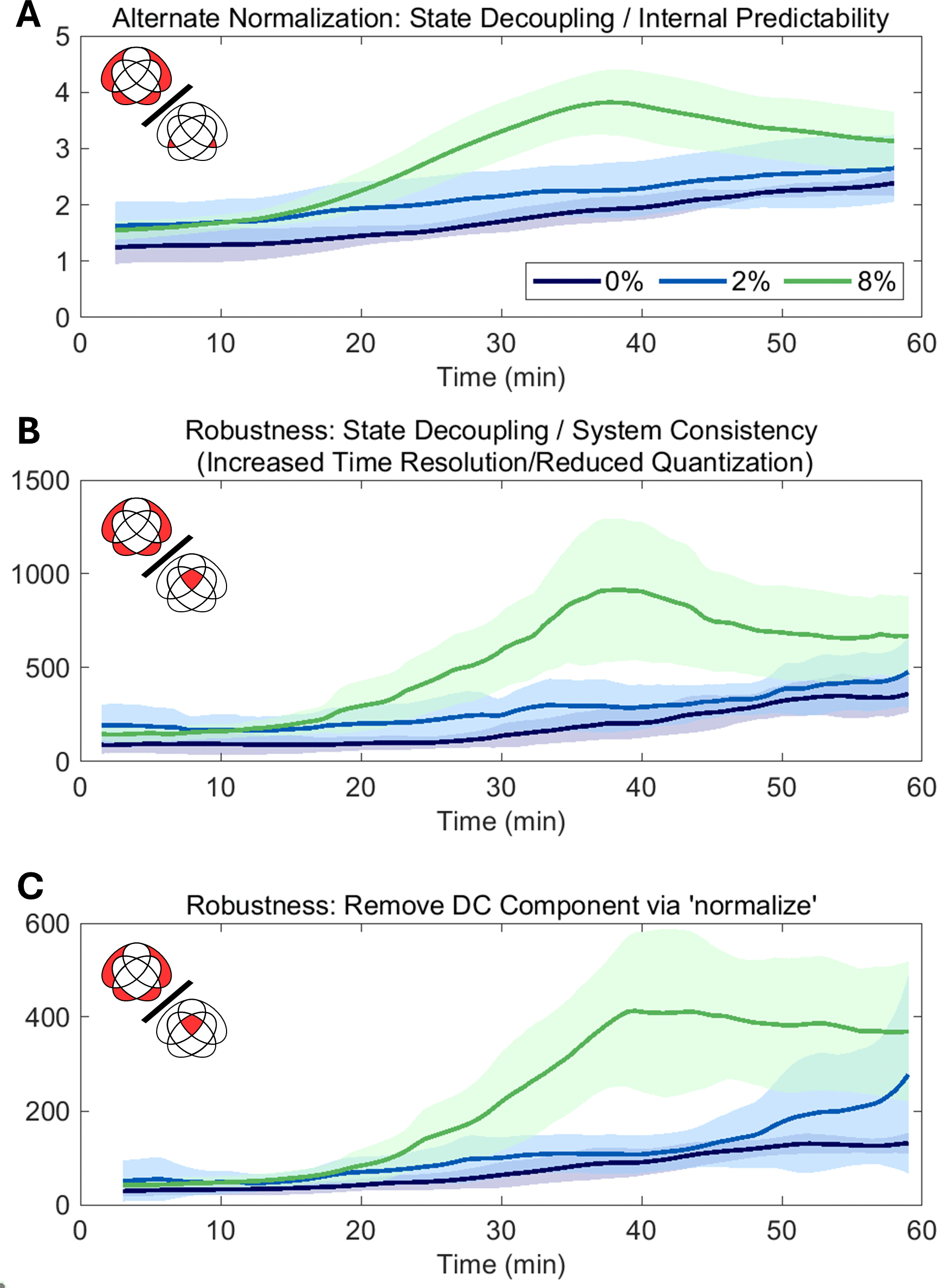
